# Prokaryotic genome design is ruled by the mechanical properties and complexity of the DNA sequence

**DOI:** 10.1101/2022.03.10.483863

**Authors:** Tsuneyuki Tatsuke

## Abstract

Although we represent the genetic code using four letters, namely A, T, G, and C, it is a chemical entity with sequence-dependent physical properties and a spatial extent. In other words, it can be said that genetic information is compressed and recorded in the genomic DNA as a storage medium with a spatial spread. This study analyzed prokaryotic genomes in terms of their physical properties and complexity. The findings clearly indicated that a mathematic equation can be used to express the relationship between the length of a genome and the upper limit of its rigidity. Furthermore, many prokaryotes have been shown to maximize DNA complexity under various physical constraints. Here, this relationship between rigidity and complexity was found to be shaped like an entropy function. It was also clarified that codon usage tendencies are affected by the physical properties and complexity of the DNA sequence. Moreover, the analysis of codons encoding two consecutive synonymous amino acids suggests that many prokaryotic codon combinations tend to increase genomic DNA complexity. This study revealed the macro-level rules for packing genomic DNA into prokaryotic cells.

## 1. Introduction

DNA sequences are usually represented with four letters: A, T, G, and C. The actual genomic DNA is, however, a heterogeneous chain with a double-helical structure consisting of four types of chemical components. The genome is, therefore, an information storage medium with a spatial extent and distinct physical properties^1^. The sequence-dependent properties of double-helical DNA have been extensively studied. At the simplest level, the sequence–structure relationships of dinucleotides as the fundamental building blocks of DNA polynucleotides have been reported^2^. There have also been reports on the sequence-dependent properties of tetranucleotide-based sequences, and the structure prediction models for long DNA sequences containing either are in good agreement with the X-ray crystal structure data of DNA^3^. Furthermore, *in silico* genomic DNA structure prediction of whole-genome folding in yeast based on these prediction models and the DNA-encoded nucleosome organization reported by Kaplan et al. (2009)^4^ reflects the actual structure of genomic DNA including the chromatin structure^1^. Therefore, genomic DNA records structural information in addition to genetic information, and it can be said that genetic information and the mechanical properties of genomic DNA are inextricably linked.

In the last decade, there have been reports of the activation of a prokaryote from chemically synthesized genomic DNA^5^, the creation of artificial bacteria with minimal genomic DNA^6^, and the generation of *Escherichia coli* bearing recoded genome using only 61 codons by synonymous codon compression^7^. A fully engineered bacterial genome, however, has not yet been created because the dynamic behavior of the entire biological system is still not fully understood. A systems-level understanding of dynamic processes such as metabolism, protein interactions, RNA transcriptional control, and the multi-layered control of all these processes is required for the complete design of biological systems, and this is an essential area of research in systems biology and synthetic biology. On the other hand, the understanding of genomic DNA based on its physical properties is also required for the complete design of biological systems, and this can be analyzed by mainly assuming a static state. In eukaryotes, the state of genomic DNA within the nucleosomes in the nucleus has been revealed using the Hi-C method^8^, and it is now known that there are three-dimensional, self-interacting genomic regions, termed topologically associating domains^9^. In prokaryotes, on the other hand, approximately 85% of the genomic DNA is composed of uninterrupted protein-coding sequences, i.e., open reading frames (ORFs), without nucleosomes and other DNA-binding proteins, and their mechanical properties are expected to be firmly ruled by trinucleotides—the codons. Indeed, A-tract clusters, which are the sequence motifs introducing the most pronounced DNA curvature, have been reported to facilitate DNA packaging in the bacterial nucleoid^10^. Besides, it is necessary to pack genomic DNA together with RNA and protein in the limited space available within prokaryotic cells. That is, genomic DNA is considered to hold not only genetic information but also structural information. Therefore, understanding of the storage of prokaryotic genome-wide genetic information requires an approach based on physical ideas.

In this study, the macro-level rules for packing genomic DNA into prokaryotic cells were revealed by focusing on the mechanical properties and sequence diversity of DNA as a heterogeneous polymer. This study was mainly based on an analysis of the mechanical properties of prokaryotic genomic DNA in units of 4 base pairs (bp)^3,11^, and diversity analysis using the Shannon entropy score^12^. From the analysis of the relationship between mechanical properties and genome length using the complete genomic DNA sequences of approximately 3,000 prokaryotic organisms registered in the National Center for Biotechnology Information (NCBI) GenBank database, the formula describing the upper limit of rigidity, a mechanical property of genomic DNA, was obtained. This formula also explains the upper limit of the size of prokaryotic genomic DNA. Of course, bottom-up approaches should consider the presence of nucleoid-constituting proteins such as HU. However, since this study aimed to discover the rules governing genomic DNA at the macro-level in a top-down manner, the effects of nucleoid-constituting proteins are also encompassed within the obtained formula. The entropy score of the genomic DNA sequence complexity showed that prokaryotes tend to maximize genomic DNA complexity as much as possible under the constraints of the mechanical properties of the genome. Moreover, the present study identified mechanical properties and sequence diversity as the driving forces that correlate codon usage with genomic DNA size and species. This study has revealed that genetic information, the mechanical properties of genomic DNA, and the complexity of DNA sequences are inseparable. These results are expected to contribute toward the creation of a fully engineered bacterial genome in the future.

## 2. Materials and Methods

### 2.1. Source of complete prokaryotic genome sequences

The NCBI GenBank (https://www.ncbi.nlm.nih.gov/genbank/) and RefSeq (https://www.ncbi.nlm.nih.gov/refseq/) assemblies, July 08, 2015, updated version, were used as sources of the prokaryotic genomic sequences, including those of eubacteria and archaea. After excluding duplicates and incomplete sequences, 2,943 sequences were obtained.

### 2.2. Data processing

All data processing was performed using Python (https://www.python.org/). The SciPy library (https://www.scipy.org/) was used for regression analysis and the Matplotlib library (https://matplotlib.org/) was used for plotting.

### 2.3. Calculations of DNA rigidity

The calculations were performed with all 136 tetranucleotide step flexibility parameters based on a potential energy surface model of a tetranucleotide^3^. The flexibility of each tetranucleotide step in a DNA sequence was calculated with a sliding step length of 1 bp^11^. After the tetranucleotide step calculations, the average value of the results was determined for a whole genomic DNA sequence in each species or for two repeated codons.

### 2.4. Entropy scoring of genomic DNA sequences

In this study, Shannon entropy was used as the entropy score for the complexity and diversity of the genomic DNA of each species. The Shannon entropy, *H*(*P*), of a whole genomic DNA sequence from each species was determined from various tetranucleotides’ *Ω* obtained from the genomic DNA sequence with a sliding step length of 1 bp, based on the following formula^12^:

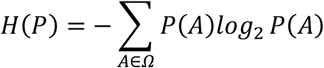

where *A* refers to the count of individual tetranucleotides, *P*(*A*) indicates the ratio of *A* in *Ω* that consists of genomic DNA sequences sliced by tetranucleotides.

### 2.5. Directionality indexes of codon pairs encoding the same amino acid

To analyze bias in the selectivity of codon pairs encoding the same two consecutive amino acids, the directionality index (DI) was used. The DI was calculated using the following formula^9^:

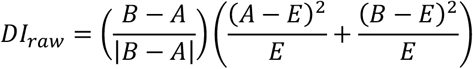

where *A* and *B* are the proportion of each codon combination in codon pairs encoding the same two consecutive amino acids, and *E* is the expected ratio of *A* and *B* under the null hypothesis and is equal to (*A* + *B*)/2. Furthermore, *B* represents the codon pair that is the reverse combination of that comprising *A*, with *B* being the more rigid of the two codon combinations. The normalized DI was defined by *DI_raw_* in the range from −1.0 to 1.0. A positive DI indicated a bias toward more rigid codon combinations, while a negative DI indicated a bias toward more flexible codon combinations. Thus, the DI values indicated both the bias strength and direction.

## 3. Results

### 3.1. The mechanical properties of DNA limit the spatial extent of prokaryotic genomes

In most prokaryotes, the protein-coding region accounts for approximately 80%–90% of the genomic DNA (Fig. 1). In other words, the codon context firmly rules most sections of the prokaryotic genome. DNA sequences are known to have mechanical properties that depend on the arrangement of the nucleotides^3^. Therefore, it was expected that prokaryotic genomic DNA, which is mostly composed of ORFs in which codons consisting of three bases are arranged regularly without interruption, would have specific mechanical characteristics. Therefore, an analysis was conducted focusing on the physical properties of the DNA including the length of the genomic DNA, using the method described in Fukue et al. (2004)^11^, and the parameters described in Packer et al. (2000)^3^ (see Materials and Methods for more details). A logistic equation type model was assumed based on the distribution shape of the upper limit of the rigidity of prokaryotic genomic DNA (Fig. 2A). Non-linear regression analysis revealed that the upper limit of the rigidity of prokaryotic genomic DNA and its size can be described by the following equation:

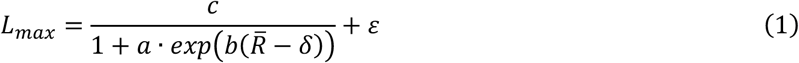

where *L_max_* indicates the maximum size of the genomic DNA, 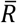 refers to the average rigidity of the genomic DNA, *ε* is a genome size of 0.058 (Mbp) of *Mycoplasma genitalium,* which is said to have the smallest genome in nature, (*δ*= 12.14316 kJ · mol^-1^ · Å^-2^) is the lower limit of the rigidity of prokaryotic genomic DNA determined from plot (Fig. 2A), and *a, b,* and *c* are the values obtained by non-linear regression analysis: *a* = 0.00854163, *b* = 2.178543, and c = 10.5789. *a* and *b* are constants, *L* = *c* + *ε* is the asymptote for the maximum theoretical genomic DNA size of prokaryotes.

**Figure 1.**
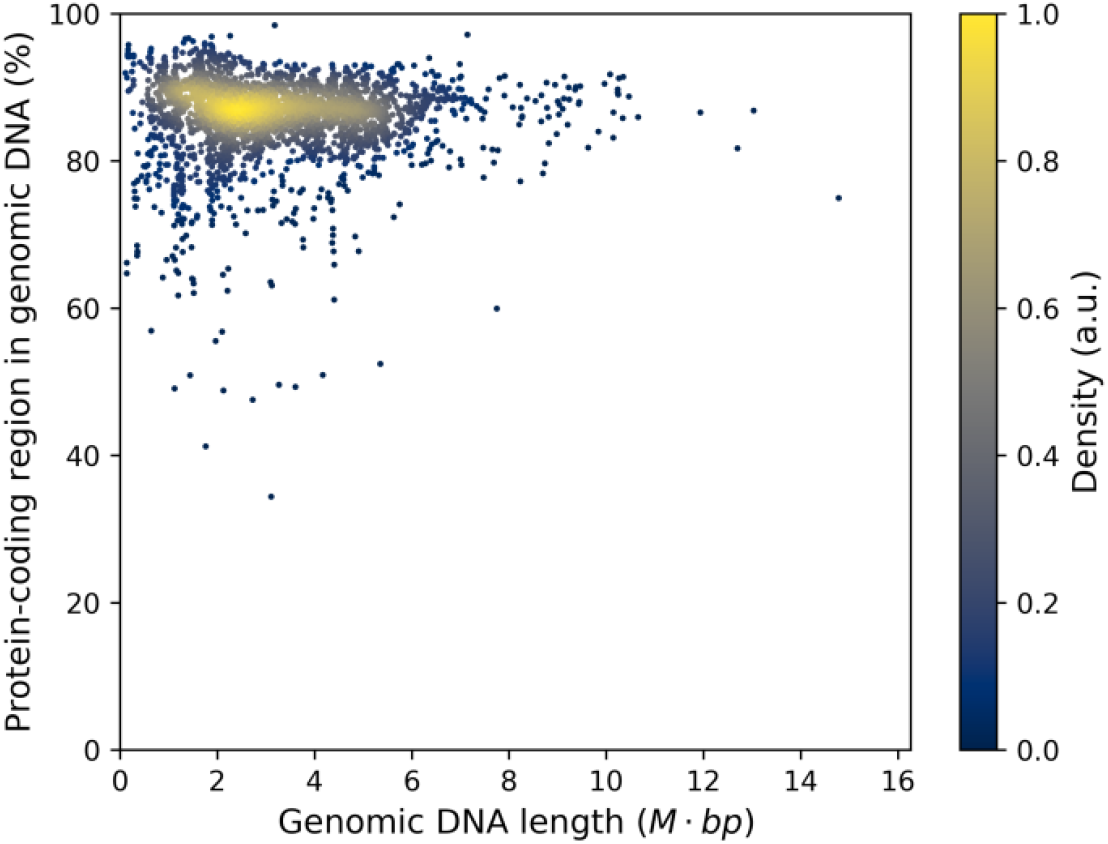
Ratios of protein-coding regions in genomic DNAs of different lengths from 2,943 prokaryotic species. Each data point corresponds to one species.

**Figure 2.**
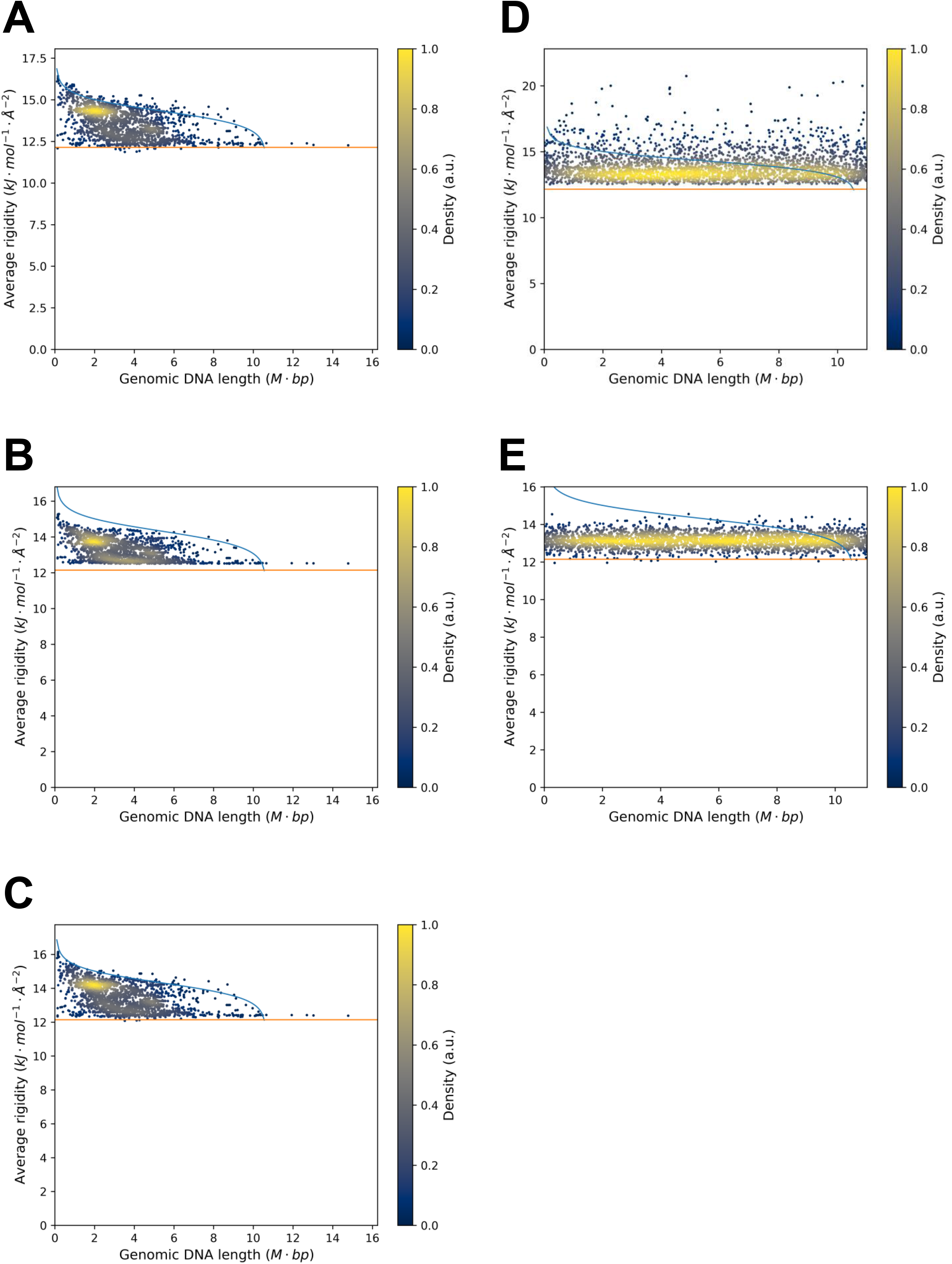
Relationship between the mechanical properties of average genomic sequence rigidity and genome length. (A) Relationship between the average rigidity and length of the genomic DNA of 2,943 prokaryotes. Each data point corresponds to one species. The blue line indicates the upper limit of the rigidity of the genomic DNA, while the orange line represents the lower limit. (B and C) Genomic DNA sequences of 2,943 prokaryotes were randomly sorted by various base unit sizes (single nucleotides, i.e., ATTGCCGAC to TGACAGCCT, or codon units, i.e., ATTGCCGAC to GACATTGCC). Genomic DNA sequences were sorted by single-base units (B), or triple-base units (C). (D and E) *In silico,* 2,943 genomic DNA sequences were randomly generated using single-base units (D), or triple-base units (E). The relationship between their rigidity and length was plotted. Each data point corresponds to one randomly generated or sorted genomic DNA sequence.

Equation (1) shows that the upper size limit of prokaryotic genomic DNA, L_max_, is ruled by its average rigidity, 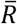. In equation (1), 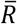 represents an average value of the energy required to deform a section of the genomic DNA per bp (kJ · mol^-1^ · Å^-2^/bp)^3^. Since the relationship between persistent length of a DNA fragment *y* (nm) and DNA rigidity *x* (kJ · mol^-1^ · Å^-2^) can be expressed by the equation *y* = 2.1246x + 10.566 [1], higher 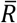 values indicate a stiffer DNA sequence and higher space costs, while lower 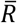 values indicate a more flexible DNA sequence and lower space costs. Contrary to expectations, the randomly sorted genomic DNA sequences of 2,943 prokaryotes by single-or triple-base units did not significantly change the distribution of species in the relationship between their rigidity and length (Fig. 2B and C), comparing to the randomly generated genomic DNA (Fig. 2D and E). Therefore, it is suggested that the set of bases that compose the genome sequences of 2,943 prokaryotes themself have the property of generating upper and lower limits that are inherent to organisms. Furthermore, such properties were not observed for genomic DNAs comprising sequences that were randomly generated with a computer (Fig. 2D and E), indicating that the relationship described by equation (1) is inherent to organisms. These findings are probably important in considering the relationship between genomic DNA length and genomic DNA folding in prokaryotes by macro-level, including effect by nucleoid-related proteins.

Solving equation (1) for the average rigidity of the genomic DNA yields:

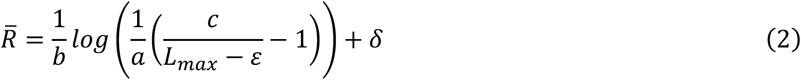

Equation (2) implies that when the size of the genome increases by a particular value, its rigidity decreases by a specific value to mitigate the space cost. In other words, prokaryotic genomic DNAs have evolved according to design rules that allow more compact storage as their size increases. The maximum theoretical size of prokaryotic genomic DNA deduced from these equations is *c* + *ε* = 10.6369 Mbp.

### 3.2. The prokaryotic genome maximizes the complexity of its nucleotide combinations under the constraints of the mechanical properties of DNA

So how does genomic DNA complexity occur under these mechanical constraints? The answer to this question lies in the diversity of the DNA sequences carried by genomic DNA. The complexity of the prokaryotic genomic DNAs was assessed based on Shannon entropy and plotted in comparison with their average rigidity. The complexity of genomic DNA and the entropy score was found to be proportional. The genomic DNA sequences of 2,943 prokaryotes produced a curve similar to the entropy function in terms of average rigidity and entropy (Fig. 3A). Further, this curve indicated that the majority of prokaryotic species lie within the curve maxima.

**Figure 3.**
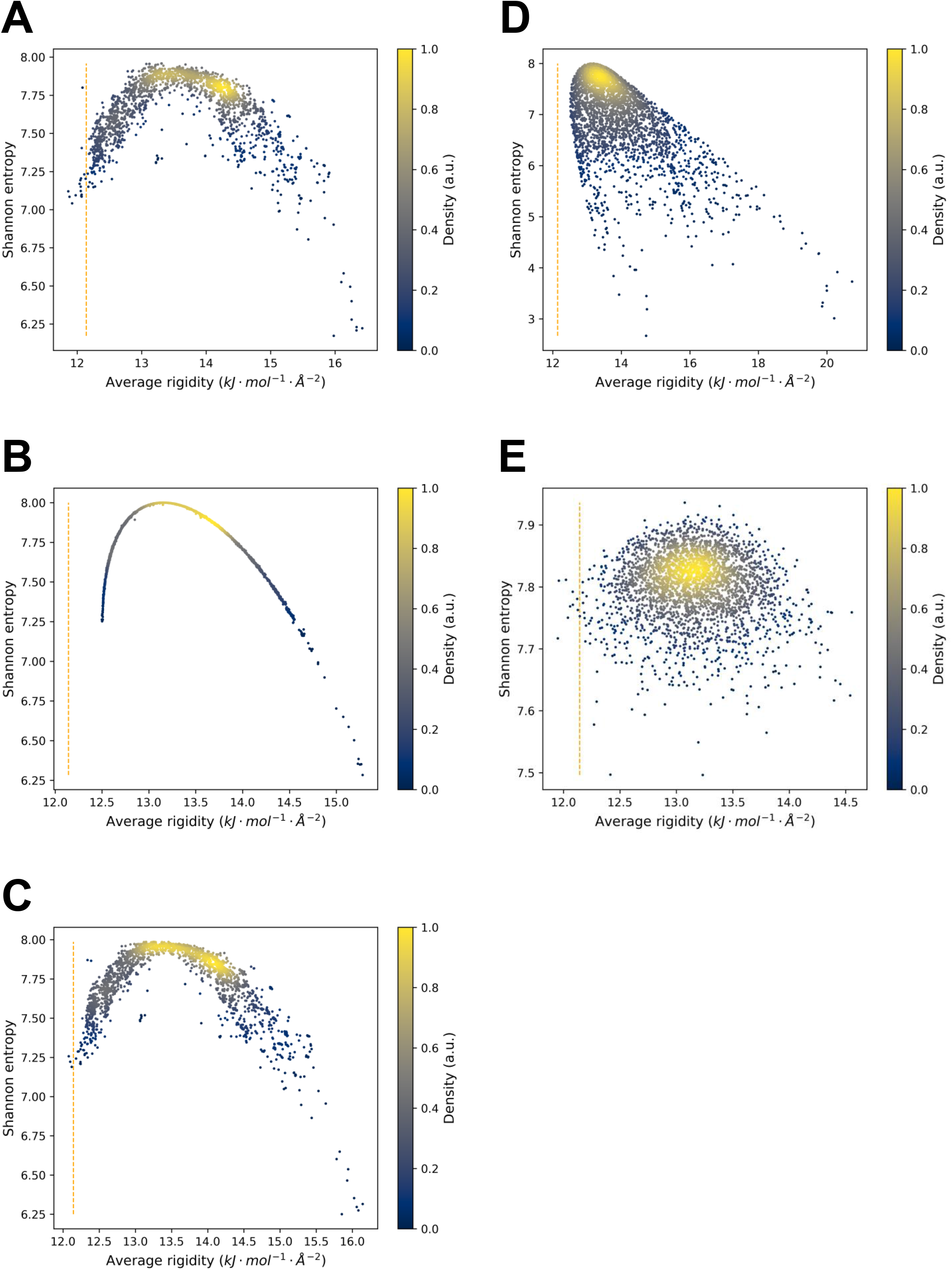
Relationship between the complexity and rigidity of genomic DNA sequences. (A) The rigidity and complexity of genome DNA sequences of 2,943 prokaryotes. Each data point corresponds to one species. (B and C) Genomic DNA sequences of 2,943 prokaryotes were randomly sorted by various base unit sizes (single nucleotides, i.e., ATTGCCGAC to TGACAGCCT, or codon units, i.e., ATTGCCGAC to GACATTGCC). Genomic DNA sequences were sorted by single-base units (B), or triple-base units (C). (D and E) *In silico*, 2,943 genomic DNA sequences were generated using various base unit sizes (single nucleotides or codons). Genomic DNA sequences were generated using single-base units (D), triple-base units (E). The relationships between their complexity and rigidity were plotted. Each data point corresponds to one randomly sorted or generated genomic DNA sequence. In all plots, the vertical axis represents Shannon entropy as a proxy for the complexity of the genomic DNA sequence, and the horizontal axis represents the average rigidity of the genomic DNA sequence. The orange dashed line indicates the lower limit of the rigidity as determined in Fig 2.

Focusing on the lower limit of the rigidity of prokaryotic genomic DNA, this value takes a fixed value of 12.14316 kJ · mol^-1^ · Å^-2^. *In silico,* 2,943 genomic DNA sequences were randomly sorted under various conditions to confirm this lower limit. When the genomic DNA sequences were sorted using units of a single base strung together randomly, the rigidity did not reach this lower limit (Fig. 3B). However, when the genomic sequence was randomly sorted using units comprising triple bases (corresponding to codons), the rigidity almost reached this lower limit (Fig. 3C). This value is considered to derive from the genomic regions consisting of three-base units, which account for approximately 85% of prokaryotic genomic DNA. In other words, the lower limit of the rigidity of prokaryotic genomic DNA is derived from the major structural unit of genomic DNA.

Comparing the plot of the genomic DNA sequences of 2,943 prokaryotes (Fig. 3A) with plots of randomly sorted genomic DNA sequences (Fig. 3B and C), the distribution shape of the plot of DNA sequences shuffled by single-base units tends to be thin and more rigid (Fig. 3B), while the distribution shape of the plot of them shuffled by triple-base units is close to that of the plot of organisms (Fig. 3C). This result is possibly due to about 85% of prokaryotic genomic DNA consists of protein-coding regions with triple-base units. The original genomic DNA shows a more diverse distribution than randomly sorted genomes, presumably because of the rules inherent to organisms for arranging DNA nucleotides. Also, comparing these plots with plots of randomly generated DNA sequences (Fig. 3D and E), it was apparent that the DNA sequences derived from organisms tend to maximize diversity as much as possible and that this is probably a property inherent in the set of bases that compose genomic DNA of organisms. In the case of genomic DNAs randomly generated by a computer and comprising single-base units, the entropy values extended downward (Fig. 3D). This indicated that these DNA sequences tended to be simpler. In contrast, when the genomic DNAs were randomly generated using triple-base units, the entropy values were biased upward (Fig. 3E).

These results suggest that prokaryotes adapt codon usage and the codon context by diversifying the genomic DNA sequence within the structures of many limitations, including mechanical limitations, environmental factors, and the size of the genetic system encoding the genomic DNA. This would help to avoid monotony in the genetic code and create genomic DNA diversity in its stead.

### 3.3. The mechanical properties and complexity of DNA sequences rule the codon context

The prokaryotic preferences for codon usage were analyzed to determine codon usage biases when encoding the same amino acid at two consecutive codons. If the genetic code functions only as a “letter,” there should be no differences in the order of codon combinations when encoding the same amino acid at two consecutive positions. However, if there is an interplay between the mechanical properties of genomes and their genetic code, it is expected that the order of such combinations would be affected. To investigate this, the directionality index (DI) was used to analyze whether the combination of codons coding the same amino acid was biased toward rigid or flexible^10^. The average rigidity of triplet repeat sequences was calculated using the method described in Fukue et al. (2004)^11^ and the parameters described in Packer et al. (2000)^3^ (see Methods for more details). Here, the Cys-Cys amino acid sequence encoded by the combination of tgc and tgt was chosen as an example of a combination that is rarely used by most species. The Ala-Ala amino acid sequence encoded by the combination of gcc and gcg was chosen as an example of a combination with different frequencies of use among species. Lastly, the Leu-Leu amino acid sequence encoded by the combination of ctt and ttg, which exerts a largely different effect on rigidity depending on the order of the codon combination, was also chosen.

The Cys-Cys amino acid sequence encoded by the combination of tgc and tgt was found to be used infrequently in either of the two consecutive codon combinations and was considered to have little structural effect on the genomic DNA. For this reason, DI bias was observed in many species (Fig. 4A). On the other hand, the Ala-Ala amino acid sequence encoded by the combination of gcc and gcg is frequently used in many species and tended to show a less biased DI in the species in which this combination is frequently used (Fig. 4B). Also, in the species with a low frequency of use of the combination of gcc and gcg, DI is similar to Cys-Cys encoded by the combination, tgc and tgt. Because it is the nature of organisms to try to diversify their genomic DNA sequences as much as possible, these relationships between frequency of use and bias appear reasonable. In addition, in the Leu-Leu amino acid sequence with a large difference in rigidity depending on the order of the combination of ctt and ttg, it was observed that many species tended to be biased toward the flexible ttg-ctt side (Fig. 4C). When such a large difference in rigidity is observed according to the order of codon combination, it is considered that each codon combination tends to have its own unique DI pattern.

**Figure 4.**
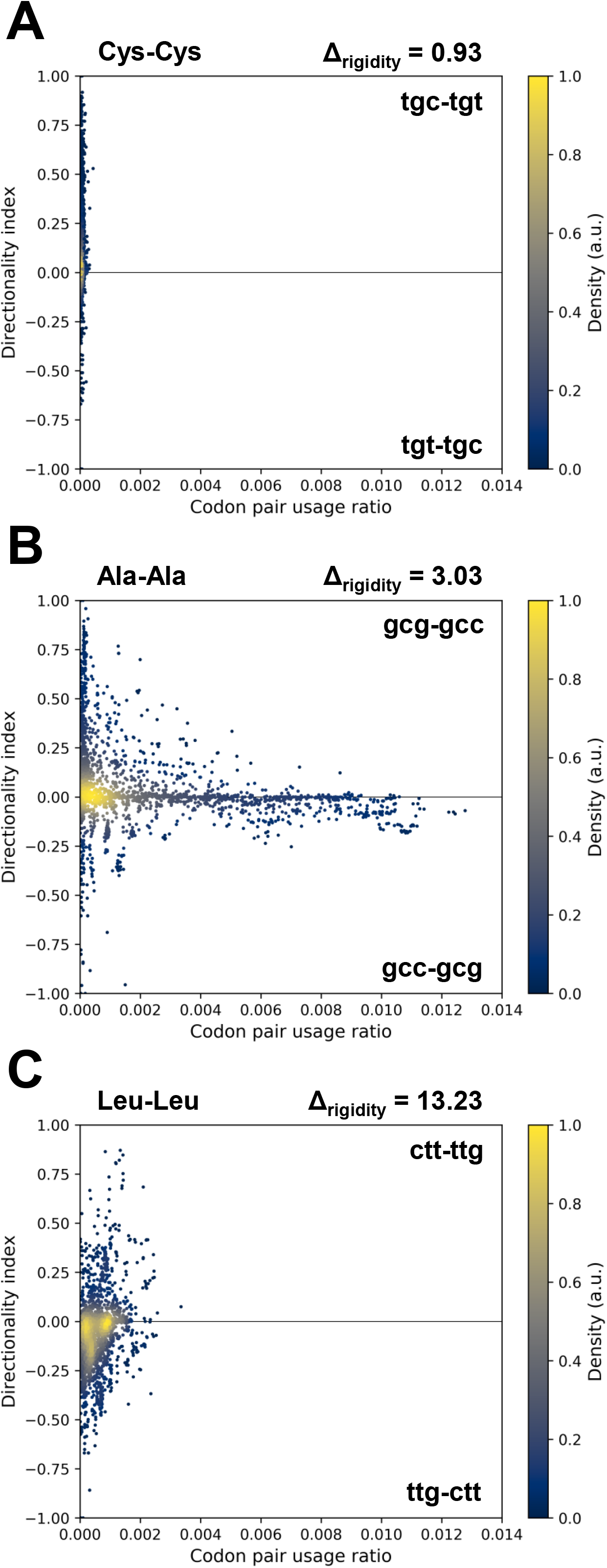
Codon usage bias when the same amino acid is encoded by two different consecutive codons. (A) When Cys-Cys is encoded by the tgc and tgt codons, the rigidity of tgc-tgt is 12.73 kJ · mol^-1^ · Å^-2^, while the rigidity of tgt-tgc is 11.80 kJ · mol^-1^ · Å^-2^. (B) When Ala-Ala is encoded by the gcg and gcc codons, the rigidity of gcg-gcc is 15.00 kJ · mol^-1^ · Å^-2^, while the rigidity of gcc-gcg is 11.97 kJ · mol^-1^ · Å^-2^. (C) When Leu-Leu is encoded by the ctt and ttg codons, the rigidity of ctt-ttg is 24.20 kJ · mol^-1^ · Å^-2^, while the rigidity of ttg-ctt is 10.97 kJ ? mol^-1^ · Å^-2^. For all plots, the directional index (DI) represents bias toward the usage of a particular combination of codons. DI takes a value from −1 to 1, with highly rigid codon combinations having a positive DI and more flexible codon combinations having a negative DI. Each data point corresponds to one species. The vertical axis represents the DI, and the horizontal axis represents the ratio of the usage frequency of the codons encoding the two target consecutive amino acids to that of all two-codon combinations. Δrigidity represents the difference in rigidity caused by the differing order of the codons encoding the same two consecutive amino acids.

These results suggest that sequences with a high usage frequency of specific codon combinations are less biased in their directionality of use, while those with a low usage frequency tend to be directionally biased toward one of the two possible combinations. This strategy appears reasonable because if the usage frequency of two codon combinations is biased, the entropy is low, but if both are used in equal proportions, the entropy is maximized. Moreover, when there is a large difference in rigidity, there is a corresponding difference observed in the distribution. The opposite tendency may be observed when the difference in rigidity is small. In most cases, however, the frequency of use of these two-codon combinations encoding the same amino acid was low relative to the total frequency of all two-codon combinations, so it is possible that the structure of the genomic DNA as a whole is not significantly affected by these combinations. It is expected that similar relationships would be observed for codon combinations encoding other amino acids, as the genetic code and its physical properties are both sides of the same coin.

## 4. Discussion

Prokaryotic genomic DNA is packed within the limited space of prokaryotic cells, and the amount of space available for packing prokaryotic genomes is difficult to expand from the viewpoint of the biochemical reactions occurring within, but it is often necessary to increase the genome size from the viewpoint of evolution. Besides, since 80%–90% of prokaryotic genomic DNA comprises protein-coding regions using three-base units, it is considered likely that the arrangement of genetic codes—the codon context—and the physical properties of genomic DNA have coevolved. The findings of this study indicate that even when different codons encode two synonymous amino acids, a bias in usage frequency is observed due to the differences in rigidity arising from the different codon combinations. These results suggest that the genetic code avoids simplifying codon combinations and evolves in concert with mechanical properties such as DNA rigidity, all while maintaining complexity.

From the above hypothesis, it would appear that prokaryotic genomic DNA maintains the evolutionary robustness of an organism by diversifying the combination of genetic codes under the limitations of restricted storage space in prokaryotic cells. It is possible that the evolution of tRNAs is also significantly affected by this aspect. In cases where codon combinations are affected by the storage space afforded to genomic DNA, the evolution of tRNAs as the decoder of codons may also be affected by the physical properties of the genetic code. It is assumed that such characteristics are driving forces in producing differences in codon usage among species.

Furthermore, the differences in genomic DNA size among prokaryotic species are due to differences in the total number of genes encoded by the genomic DNA. In other words, differences in the quantity of genetic information and the structure thereof create the observed species differences in genomic DNA. These differences could be the result of the environmental adaptation and diversification of prokaryotes. Some prokaryotes have a larger genome than *Saccharomyces cerevisiae* (1.2 × 10^7^ bp), and the average rigidity of their genomic DNAs is around the lower limit of the rigidity of prokaryotic genomic DNA determined here (shown in Fig. 2A). From this perspective, it is possible that eukaryotes have acquired introns and exons to overcome the limitations exerted on codon placement strategies by the mechanical constraints evident in prokaryotic genomes. It is possible that eukaryotic genomic DNA was released from these mechanical restraints by dividing the genome into ORFs, and even further diversification was made possible by acquiring nucleosomes that store genomic DNA efficiently. Therefore, the rigidity of genomic DNA may have been one of the driving forces in the evolution from prokaryote to eukaryote.

In recent years, researchers in synthetic biology have synthesized prokaryotic total genomic DNAs^5-7^. However, full, made-from-scratch prokaryotic genomes have not yet been designed. The findings of the current study should facilitate the future design of synthetic whole prokaryotic genomes.

## Acknowledgments

I thank Dr. Hiroyuki Hamada (Kyushu University) for advice on mathematical model construction and other aspects of the study and his insights regarding the manuscript content. I thank Natasha Beeton-Kempen, Ph.D., from Edanz Group (https://en-author-services.edanzgroup.com/) forediting a draft of this manuscript.

